# Precision size and refractive index analysis of weakly scattering nanoparticles in polydispersions

**DOI:** 10.1101/2021.11.13.468485

**Authors:** Anna D. Kashkanova, Martin Blessing, André Gemeinhardt, Didier Soulat, Vahid Sandoghdar

**Affiliations:** Max Planck Institute for the Science of Light, 91058 Erlangen, Germany; Max-Planck-Zentrum für Physik und Medizin, 91058 Erlangen, Germany; Department of Physics, Friedrich-Alexander University Erlangen-Nürnberg, 91058 Erlangen, Germany; Department of Clinical Microbiology, Immunology and Hygiene, Friedrich-Alexander University Erlangen-Nürnberg and Universitätsklinikum Erlangen, 91054 Erlangen, Germany

## Abstract

Characterization of the size and material properties of particles in liquid suspensions is in very high demand, e.g., for the analysis of colloidal samples or of bodily fluids such as urine or blood plasma. However, the existing methods are limited in deciphering the constituents of realistic samples. Here, we introduce iNTA as a new method, which combines interferometric detection of scattering with nanoparticle tracking analysis, to reach an unprecedented sensitivity and precision in determining the size and refractive index distributions of nanoparticles in suspensions. After benchmarking iNTA with samples of colloidal gold, we present its remarkable ability to resolve the constituents of various multi-component and polydisperse samples of known origin. Furthermore, we showcase the method by elucidating the refractive index and size distributions of extracellular vesicles from *Leishmania* parasites and nanoparticles in human urine. The current performance of iNTA already enables advances in several important applications, but we also discuss possible improvements.

---

Size and refractive index represent two key attributes of nanoparticles that are found in a wide range of disciplines such as medicine,^1^ pharmaceuticals,^2^ food industry,^3^ and agriculture.^4^ The samples of interest often consist of natural or synthetic suspensions of different origin and make. For example, bodily fluids such as blood plasma, cerebro-spinal fluids, or urine contain bioparticles such as extracellular vesicles, covering a large spectrum of size and protein or RNA content, which serve as disease marker.^5,6^ Information about the size, material and abundance of particles in such heterogeneous mixtures is highly desirable in fundamental research as well as clinical and industrial applications.

Various techniques can be employed to determine the particle size distribution.^7^ Electron microscopy (EM) provides an exquisite resolution in direct imaging, but its sample preparation procedure, low speed and ex-situ operation strongly hamper its appeal. Indeed, optical methods dominate the diagnostic and analytical arena despite their intrinsic diffraction-limited resolution because they are fast and can be applied to a broad set of samples in the liquid phase. One of the workhorses is dynamic light scattering (DLS), which makes use of temporal correlations in the light scattered by an ensemble of diffusing particles.^8^ In this method, the particle size is extracted from the statistical analysis of the autocorrelation function of the light intensity. Today, DLS is the most commonly used technique in particle sizing, as it is easy to use and offers high accuracy for averaged values. However, this method has a low size resolution, thus, confronting limits in the analysis of polydisperse solutions.^8^ A more recent approach, referred to as nanoparticle tracking analysis (NTA), analyzes the trajectories of individual particles to quantify their diffusion constants (*D*) and, thus, diameter *d* (for simplicity we consider spherical nanoparticles throughout this work, see also Eq. 1 below).^9,10^ Conventional NTA employs dark-field microscopy in which the signal is proportional to the scattering cross section (*σ*_sca_) of a particle and, thus, scales as *d*^6^. This rapidly lowers the sensitivity of NTA for smaller particles. Currently, leading NTA instruments are validated for gold nanoparticles (GNP) as small as 30 nm and polystyrene (PS) particles larger than 60 nm.^10,11^ Furthermore, holography has been used for imaging and tracking particles, but the reported sensitivity corresponds to the scattering strength of PS particles with *d ∼* 300 nm.^12–15^ Thus, there is a need for methods with higher sensitivity in order to access more weakly scattering particles and for better resolution in deciphering the constituents of heterogeneous mixtures. In accomplishing the latter goal, it is also very helpful to obtain valuable insight about the material make of the particles under study. Indeed, the scattering signal in NTA has been used to assess the refractive index, but the precision in these studies has also been strongly affected by the limited signal-to-noise ratio (SNR) in dark-field microscopy.

In this work, we introduce iNTA as a new method that employs interferometric detection of scattering to analyze the trajectories and scattering cross sections of diffusing single nanoparticles. Interferometric detection of scattering, coined iSCAT,^16,17^ offers a highly efficient optical contrast, which has been used by a range of methods for label-free sensing of single proteins,^18^ mass photometry^19^ and high-speed tracking of transmembrane proteins.^20^ Here, we exploit the high SNR of iSCAT to achieve a high precision in determining *D*, and thus the particle size *d*. In addition, we perform quantitative measurements of the iSCAT contrast to assess the scattering cross section of each nanoparticle, providing direct information about its refractive index. We present an unprecedented precision and resolution in measuring the size and the refractive index of nanoparticles, not only in monodisperse, but also in polydisperse mixtures in the range of *d ∼* 10 *−* 200 nm and of complex entities such as layered particles. Furthermore, we exhibit the advantages of iNTA in exemplary field applications such as the characterization of extracellular vesicles from parasites and human urine.

## Results

### Measurement principle

The diffusion of a particle in a fluid is described by the Stokes-Einstein (SE) equation

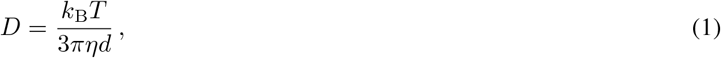

where *k*_B_ is the Boltzmann constant, *T* and *η* are the temperature and viscosity of the fluid, respectively, and *d* signifies the (apparent) diameter of the particle.^21^ Thus, one can arrive at *d* by evaluating *D* from the mean squared displacement (MSD) of a particle trajectory. Because the number of trajectory points affects the measurement precision, fast recordings are highly desirable. However, high-speed imaging can only help if a large SNR is maintained to ensure low localization error.^22^ This is where iSCAT microscopy provides a decisive advantage due to its ability to track nanoparticles with a high spatial precision and temporal resolution.^17^

Another quantity of importance in our work is the scattering cross section *σ*_sca_ of a nanoparticle. For uniform Rayleigh particles with *kd «* 1, where *k* is the wavenumber, *σ*_sca_ *∝* |*α*|^2^, where

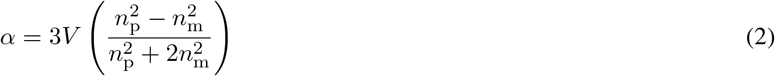

represents the polarizability.^23^ Here, *V ∝ d*^3^ denotes the particle volume, and *n*_p_ and *n*_m_ are the refractive indices of the particle and its surrounding medium, respectively. The recorded iSCAT signal is proportional to the electric field of the scattered light and, thus, to *α*.^16^ It is expressed as a contrast (*C*) and can be read from the central iPSF lobe. For particles with *kd ≳* 1 and for multi-layered particles, a generalized Mie theory describes the scattering strength (see Supplementary Information (SI)). As we shall see, the information about *C* plays a decisive role in deciphering various species and determining their refractive indices in a polydispersion.

Figure 1a sketches a common wide-field setup for performing iSCAT measurements.^17^ In the left column of Fig. 1b, we display three examples of the interferometric point-spread function (iPSF) that results from the interference of planar (reflected from the sample interface) and spherical (scattered by the particle) waves.^17,24^ The iPSFs vary qualitatively depending on the particle position relative to the coverslip and the focal plane.^24^ To localize an iPSF in a given image, we apply radial variance transform (RVT), which converts the iPSF into a bright spot,^25^ as shown in the right column of Fig. 1b. An example of a trajectory is overlaid in Fig. 1c.

**Figure 1:**
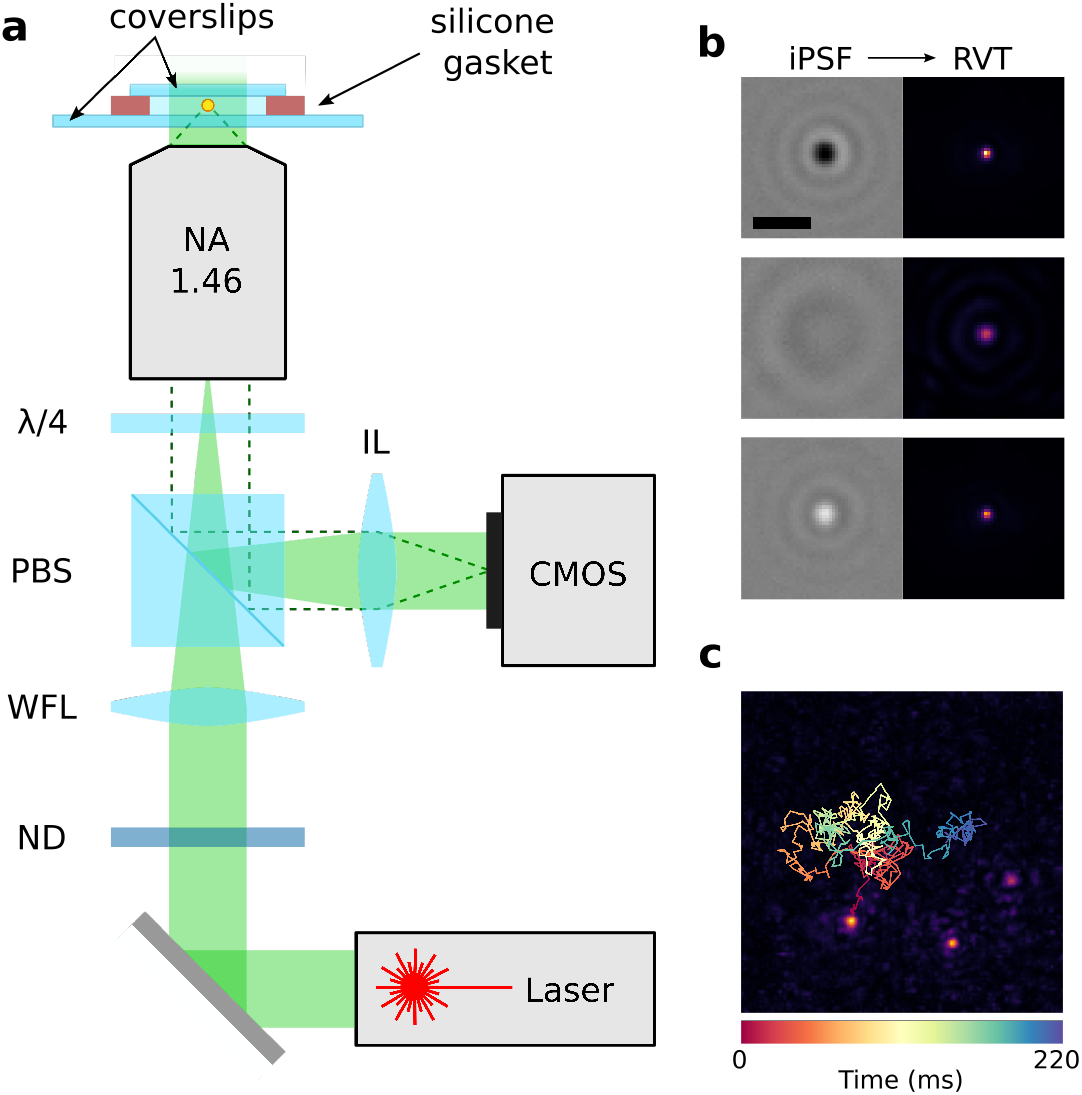
iSCAT setup and trajectory extraction. **(a)** Wide-field iSCAT setup for tracking freely diffusing particles. Linearly polarized light from a laser traverses a neutral density filter (ND) used to adjust the incident power, passes a polarizing beam splitter (PBS) followed by a *λ/*4 plate that renders its polarisation circular. A wide-field lens (WFL, *f* = 400 mm) focuses the light at the back focal plane of the objective. An imaging lens (IL, *f* = 500 mm) projects the reflected (solid green area) and scattered (dashed line) light on a CMOS camera chip. **(b)** Three examples of iPSF (left) and the corresponding RVT images (right). Scale bar is 1 µm. **(c)** A 7 × 7 µm^2^ frame showing three 15 nm GNPs with an overlaid trajectory of one GNP recorded over 220 ms.

We introduce a dilute suspension of nanoparticles in a closed chamber on a microscope coverslip (see Fig. 1a) and image the diffusing entities using a fast camera. The focal plane of the microscope objective is typically placed at a few micrometers above the coverslip and is stabilized with an active focus lock. The trajectory lengths are predominantly limited by the axial diffusion of particles.

We extract *D* and thereby *d* by fitting the MSD plot for individual trajectories. For monodisperse samples, we can also evaluate the mean diffusion constant 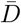 as well as a localization error by fitting averaged MSD plots, weighted by the trajectory length (Fig. 2a). For polydisperse samples, we additionally exploit the knowledge of *C*. Here, we report the maximum positive contrast from each trajectory because the interferometric contrast modulates in the axial direction as the particle traverses the illumination area.^17^ We note that the systematic and technical errors encountered in our measurements are common to all NTA experiments and stem from uncertainties in temperature, drift, and vibrations.^26^ Furthermore, it is worth noting that the common-path nature of our interferometric measurements makes them very robust against spurious phenomena that might affect the optical path.

**Figure 2:**
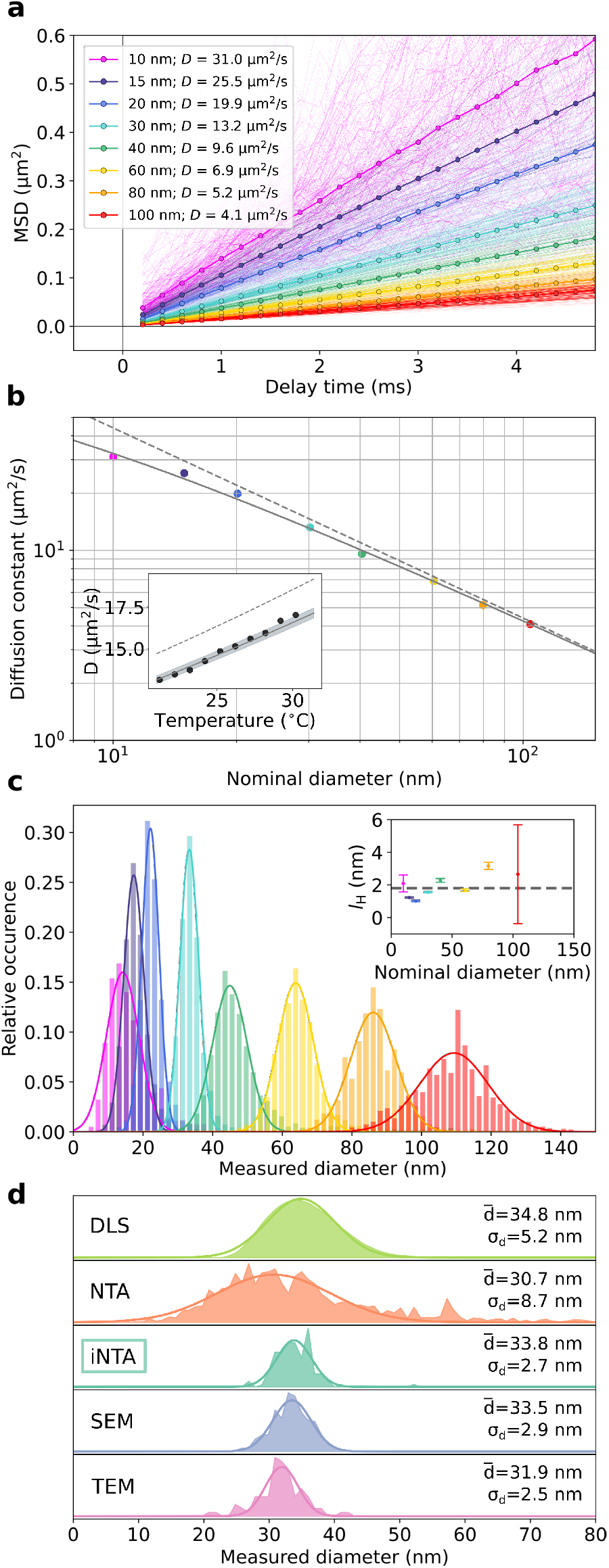
Monodisperse particle samples. **(a)** Mean square displacement (MSD) versus delay time for GNP samples of different sizes. Thin lines show the MSD extracted from each individual trajectory. Thick lines display the weighted average (by trajectory length). Diffusion constants extracted from the fits are listed in the legend. **(b)** Diffusion constants extracted from the data in (a) versus the nominal GNP diameter provided by the manufacturer. Dashed grey line indicates the SE relation for *T* = 21°C. Solid line is a fit to the SE relation with an offset in particle radius by the hydration layer thickness. Inset: Diffusion constants for 30 nm GNPs at different temperatures. Solid line and shaded area indicate the calculated diffusion constant considering *d*_H_ based on a hydration layer of *l*_H_ = 1.8 ± 0.3 nm. Dashed line shows the outcome of SE for *d*_nom_. **(c)**. Histograms of particle diameters extracted from the SE relation. Individual measurements were weighed by their trajectory lengths. The data for 10, 15, 20 and 30 nm GNPs were recorded at 40 mW illumination power; the rest was recorded at 2 mW. Inset: symbols show *l*_H_ and its error bar. Dashed line indicates the value of *l*_H_ obtained from the global fit in (b). **(d)** Comparison between different measurement techniques. The output of DLS measurements represents the intensity-weighted distribution.

### Monodisperse particle samples

We start by applying iNTA to commercially available monodisperse samples in order to benchmark its performance. Figure 2 summarizes the outcome of our measurements on GNPs from two different manufacturers. The thin lines in Fig. 2a display MSD curves from individual trajectories that contained at least 25 localization events. The thick curves show the resulting linear averaged MSD plots, which confirm free diffusion.

Figure 2b displays the measured mean values of *D* for GNPs of various diameters. The dashed line presents the behavior of the SE relation expected for the nominal particle diameters (*d*_nom_) provided by the manufacturer. Although the agreement with the data is satisfactory (note the logarithmic scales), the high precision of our measurements reveals small deviations, which suggest a systematic correction to the particle size. Indeed, the solid curve in Fig. 2b reports an excellent agreement between theory and experiment if we consider an increase of the radius by *l*_H_ = 1.8 *±* 0.3 nm for all particles. We attribute the main effect to a hydration (stagnant) layer,^27^ but additional contributions might also come from surfactant molecules. To examine the SE relation further, we also performed measurements at different temperatures using a micro heating stage (VAHEAT, Interherence). As exemplified for the case of 30 nm GNPs in the inset of Fig. 2b, we find a very good agreement between the experimental data (symbols) and the prediction of Eq. (1) when replacing *d*_nom_ (dashed line) by a hydrodynamic diameter *d*_H_ = *d*_nom_ + 2*l*_H_ (solid line).

In Fig. 2c, we display the histograms of the measured diameters (*d*_mes_) extracted from individual GNP trajectories. Gaussian fits to the data establish normal distributions, allowing us to determine a mean value 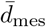 and a standard deviation 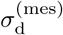.^28^ Table 1 presents these data as well as the uncertainty (Δ*d*_mes_) in determining 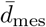. In addition, we list the extracted values of 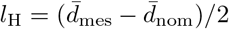 and its error bars (Δ*l*_H_) for each measurement series (see also inset in Fig. 2c). We verified that the measured quantities do not depend on the incident laser power, camera chip illumination, and the focal plane position.

**Table 1:** Nominal and measured properties of various GNP samples marked by sub- or superscript labels “nom” and “mes”, respectively. All quantities are in units of nm. See text for the definitions of 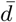and *σ*_d_. Δ*d* is computed as the 95% confidence interval of the mean by multiplying the standard error of the fit by 1.96.^28^ 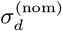 is calculated from the manufacturer (BBI Solutions, Sigma Aldrich) specified coefficient of variation. *l*_H_ represents the thickness of the hydration layer determined as (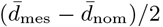). Δ*l*_H_ is the error on the hydration layer thickness calculated as 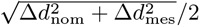. The number of extracted trajectories as well as the total number of localizations (in millions) are indicated.

| Manufacturer | $\bar{d}_{\text{nom}}$ | $\Delta d_{\text{nom}}$ | $\sigma_d^{(\text{nom})}$ | $\bar{d}_{\text{mes}}$ | $\Delta d_{\text{mes}}$ | $\sigma_d^{(\text{mes})}$ | $l_{\text{H}}$ | $\Delta l_{\text{H}}$ | Traj. | Locs. ($\times 10^6$) |
| --- | --- | --- | --- | --- | --- | --- | --- | --- | --- | --- |
| BBi | 10 | 1 | $\leq 1$ | 14.2 | 0.3 | 4.7 | 2.1 | 0.6 | 819 | 0.04 |
| BBi | 14.9 | — | $\leq 1.5$ | 17.4 | 0.1 | 2.9 | 1.3 | 0.0 | 11341 | 1.28 |
| BBi | 20.1 | — | $\leq 1.6$ | 22.1 | 0.1 | 2.5 | 1.0 | 0.0 | 6635 | 1.35 |
| BBi | 30.2 | — | $\leq 2.4$ | 33.2 | 0.1 | 2.6 | 1.6 | 0.1 | 1535 | 0.83 |
| BBi | 40.4 | — | $\leq 3.2$ | 45.0 | 0.2 | 5.0 | 2.3 | 0.1 | 2943 | 1.24 |
| BBi | 60.5 | — | $\leq 4.8$ | 63.9 | 0.2 | 5.1 | 1.7 | 0.1 | 2401 | 1.48 |
| BBi | 79.8 | — | $\leq 6.4$ | 85.9 | 0.4 | 6.6 | 3.2 | 0.2 | 1234 | 1.15 |
| SA | 104 | 6 | $\leq 8$ | 109.3 | 0.9 | 9.8 | 2.7 | 3.0 | 712 | 0.97 |

To compare iNTA with the existing state-of-the-art methods, we used DLS (ZetaSizer ZS90), NTA (Nanosight NS500), Scanning electron microscopy (SEM, Hitachi S-4800) and transmission electron microscopy (TEM, Zeiss EM10) instruments to characterize GNPs with a nominal diameter of 30 nm, as an exemplary sample at the lower limit of the commercial NTA. The results of these measurements are summarized in Fig. 2d. It is evident that the DLS and NTA size distributions have larger spreads than those of SEM and TEM measurements. The width of the iNTA distribution, however, rivals that of TEM, thus, combining an excellent *resolution* with the advantages of optical measurements. We note that the *accuracy* of each method depends strongly on careful calibrations and consideration of various systematic effects.

### Polydisperse particle samples

One of the central demands on sensing and sizing technologies is the identification of different species in a mixture. To investigate the performance of iNTA in such applications, we prepared various mixtures. First, we considered a suspension containing 15 nm, 20 nm and 30 nm GNPs. To set the ground, in Fig. 3a, we show the intensity-weighted distribution of a DLS measurement (ZetaSizer ZS90), yielding a continuous featureless distribution. As displayed in Fig. 3b, conventional NTA (Nanosight NS500) does not resolve the different populations either. In this case, we have also plotted a histogram of the scattering intensities along the right-hand vertical axis.

**Figure 3:**
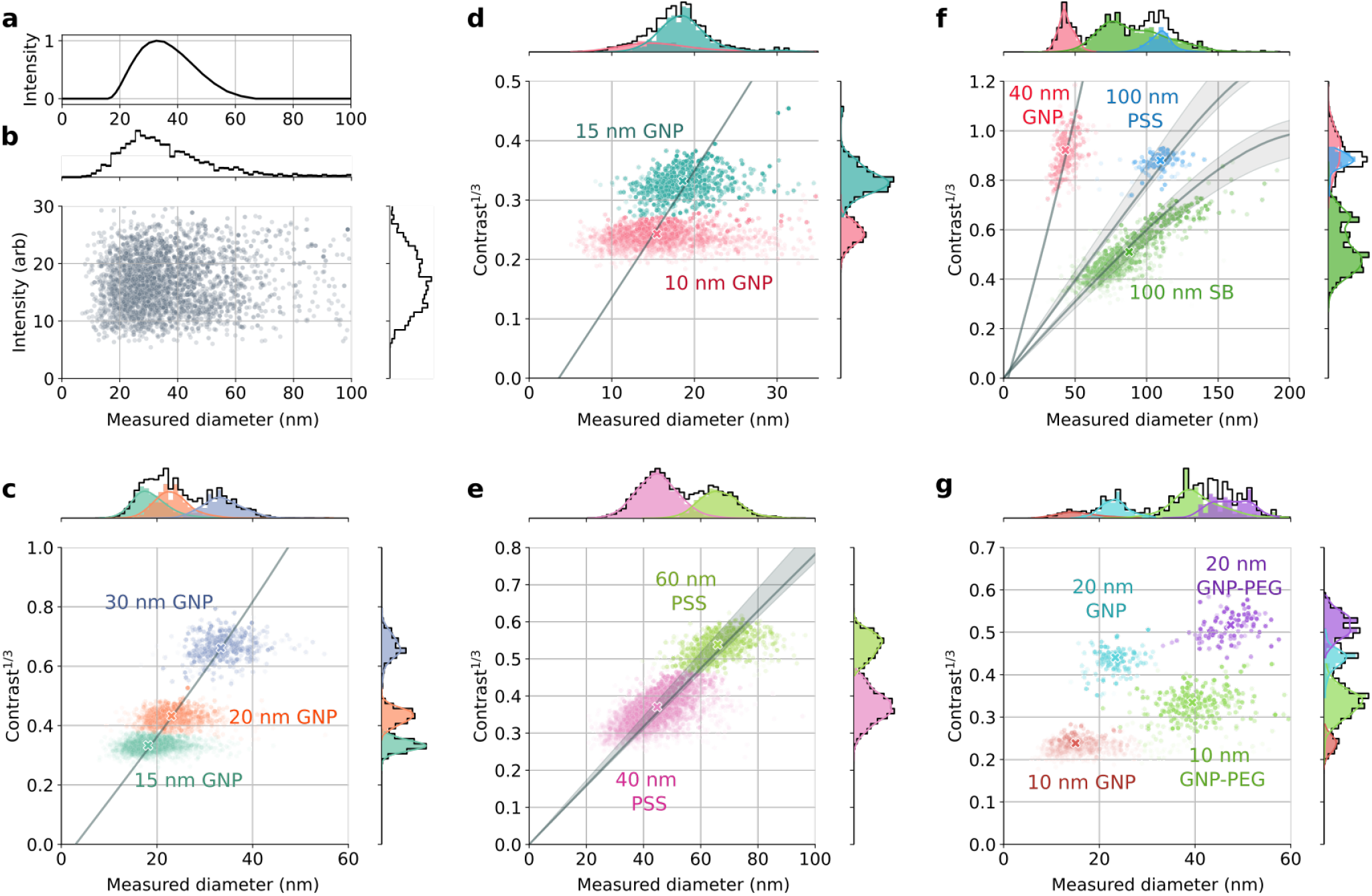
Polydisperse particle samples. DLS **(a)**, NTA **(b)**, and iNTA **(c)** measurements of a mixture of 15, 20 and 30 nm GNPs. **(d-g)** iNTA measurements of various mixtures as labeled in each graph. The horizontal and vertical axes denote the measured diameter and the third root of the iSCAT contrast, respectively. The transparency of each data point indicates the length of its trajectory. In each figure, a 2D Gaussian mixture model is used to identify different populations highlighted in color. The grey curves establish the relationship between 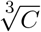 and *d*_mes_ according to the respective refractive indices while the shaded regions indicate the uncertainties in the refractive index data. Crosses in (c-g) signify the medians of each data cloud.

The scatter plot in Fig. 3c shows 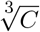 and *d*_mes_ for individual trajectories extracted with iNTA. Motivated by the fact that *C ∝ d*^3^, the choice of 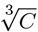 conveniently provides a dimension on a par with *d*, and its significance is in representing the scattering strength which is in turn related to the refractive index (see Eq. (2)). A visual inspection of the data clearly reveals three clusters. In fact, the histogram of 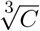 values plotted on the right-hand vertical axis also resolves the three populations on its own. Application of a two-dimensional (2D) Gaussian mixture model (GMM) with full covariance^29^ lets us decipher the three populations in a quantitative manner and identify the populations in the *d*_mes_ histogram. In Figs. 3d and 3e, we show other examples, where iNTA fully resolves mixtures of 10 nm and 15 nm GNPs and of 40 nm and 60 nm PS spheres (PSS), respectively even though particles in this size range are usually not accessible in methods based on dark-field microscopy.^30^

The advantage of combining the knowledges of *C* and *d*_mes_ becomes even more apparent when particles of similar size or *σ*_sca_ are analyzed, as shown in Fig. 3f for a mixture of 40 nm GNPs, 100 nm PSS and 100 nm silica beads (SB). Neither *C* nor *d*_mes_ alone can provide clear information about the composition of the sample, but their combination in a 2D scatter plot provides a very robust evidence for the existence of three different species. Again, as shown by the color-coded overlays, application of a GMM analysis allows us to decompose the *C* and *d*_mes_ histograms. We note in passing that the horizontal stretch of the data clouds in Figs. 3c and 3d are due to the uncertainty resulting from short trajectories or localization errors. The diagonal extension of the data in Figs. 3e and 3f, however, reveals the true size distribution in the sample.

Measurements of *C* and *d*_mes_ also provide direct access to the refractive index (RI) of the particles. Here, we considered RI for gold (*n*_Au_ = 0.63 + 2.07*i*)^31^ and fitted the data in Figs. 3c, 3d and 3f, resulting in the solid lines. The intercept of the horizontal axis yields another independent measure for the hydration shell 2*l*_H_, which amounts to 1.6 nm, 1.8 nm, and 1.5 nm for the three cases respectively. We used this information to relate the experimentally measured *C* and the expected value of *σ*_sca_ with one single calibration parameter for our setup. Next, we use this calibration and fit the data for PSS and SB in Figs. 3e and 3f to arrive at *n*_PS_ = 1.62 and *n*_Si_ = 1.45, which are in good agreement with the literature values signifed by the grey bands.^32–34^ We remark that for PSS and SB, the RI curves calculated from the full Mie theory^23^ deviate from a straight line because *σ*_sca_ for larger particles begins to contain contributions from higher order multipoles.

We also investigated a more complex mixture of 10 nm and 20 nm GNPs with and without polyethylene glycol coatings (Creative Diagnostics; molecular weight of PEG 3,000). Figure 3g showcases the high performance of iNTA by clearly distinguishing four populations. Moreover, the measurements provide us with the direct assessment of the thickness of the PEG layer, which in this case corresponds to about 12 nm. These results pave the way for future sensitive and quantitative investigation of composite nanoparticles and their interaction with the surrounding liquid phase.^35^

The superior sensitivity and resolution of iNTA measurements on monodisperse and known polydisperse nanoparticles prompted us to employ it in realistic field problems. Indeed, there is a significant number of applications in which nanoparticles of various substances and sizes need to be characterized in a fast, accurate, and non-invasive manner. Here, we discuss the analysis of synthetically produced lipid vesicles as well as extracellular vesicles (EV), which contain various proteins, nucleic acids, or other biochemical entities either in their interior or attached to them. EVs have been identified as conveyers for cell-cell communication and as disease markers,^5,6^ but studies are partly hampered by the throughput and resolution in their quantitative assessment.^36,37^ EVs are often grouped as *exosomes* (diameter 30-150 nm, originating from inside a cell) and *microvesicles* (diameter 100-1000 nm, stemming from the cell membrane), while particles smaller than 150 nm might also be referred to as small extracellular vesicles (sEVs).^38,39^ We now discuss three case studies.

### Synthetically produced liposomes

Figure 4a shows the outcome of iNTA measurements on a sample of synthetically produced liposomes, which was filtered to exclude particles larger than about 200 nm. Liposomes consist of a lipid bilayer shell surrounding an aqueous interior (see inset in Fig. 4a) and can, therefore, be modelled by a generalized Mie theory^23^ that takes into account the thickness (*t*_sh_) and RI (*n*_sh_) of the shell as well as the RI of the interior (*n*_in_). In Fig. 4b, we plot the calculated isosurfaces of constant *C* and *d*_mes_ as a function of *t*_sh_, *n*_sh_ and *n*_in_ for three exemplary points marked in Fig. 4a. We find that all three surfaces meet when *n*_in_ ≃ 1.334 for water. Furthermore, if we assume the published value of *n*_sh_ = 1.48 for our lipids,^40^ we deduce *t*_sh_ = 5.8 nm. Indeed, the orange curve in Fig. 4a confirms that the model based on these values fits the whole experimental data extremely well. To illustrate the sensitivity of the results to variations of the liposome inner part, we also plot the dashed curves in Fig. 4a for larger *n*_in_ but the same shell parameters. Remarkably, the high SNR of iNTA lets us clearly discriminate against a simple model for a uniform nanosphere.

**Figure 4:**
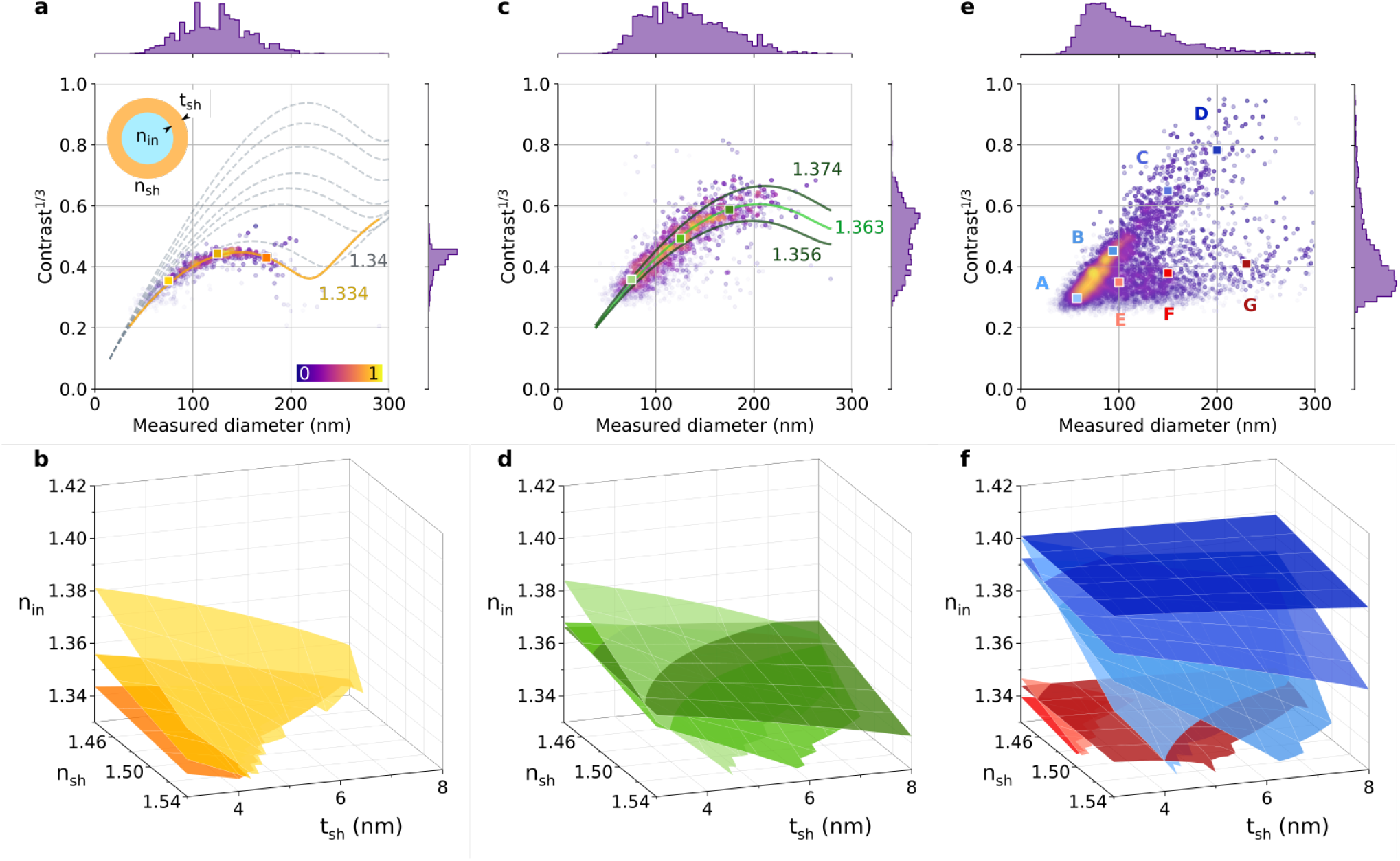
Extracellular vesicles. (**a**,**c**,**e)** iNTA scatter plots of synthetically produced liposomes (a), EVs from *Leishmania* parasites (c), and EVs from urine of a healthy human donor (e). **(a)** Color bar denotes point density. The grey dashed contours correspond to different refractive indices of the inner part of the liposomes, starting at 1.334 (water) and increasing in steps of 0.02 from 1.34. Inset shows a cartoon of a vesicle. **(c)** The green line indicates the best fit value for *n*_in_, while the olive lines indicate the 25th and 75th percentile of the extracted *n*_in_. **(b**,**d**,**f)** Isosurfaces of constant particle size and iSCAT contrast for the points marked in (a,c,e) over a range of values for *n*_in_, *n*_sh_ and *t*_sh_.

### Parasite EVs

Leishmaniasis is a potentially mortal disease, classified by the World Health Organisation (WHO) as one of the 20 neglected tropical diseases worldwide. *Leishmania* parasites secrete numerous virulence factors, most of which are carried together with small RNA inside EVs.^41,42^ Quantitative characterization of the vesicles emitted by *Leishmania* (LEVs) would be of great value for understanding their role in the infection process, but reliable data are missing.

In Fig. 4c, we present an iNTA scatter plot of LEVs. The size histogram is consistent with recently published results^43^ from DLS and NTA. The iNTA data, however, provide access to more quantitative insight. In Fig. 4d, we again use a shell model and examine the isosurfaces of three points marked in Fig. 4c. We find that *n*_in_ remains well bounded to a tight interval of (1.33, 1.38) if we allow for *t*_sh_ *∈* (3, 8) nm and *n*_sh_ *∈* (1.44, 1.54) in order to account for various lipid shell thicknesses and up to 60% protein content in the shell.^44–46^ The deduced values of *n*_in_ imply that particles of different sizes are sparsely loaded and are mostly made of water.

Extracellular vesicles are usually grouped in two classes of exosomes and microvesicles with different cellular origins.^6^ Nevertheless, the confined 2D data cloud in Fig. 4c and its smooth distribution suggest that all observed LEVs have a similar composition. To elucidate this further, we display the green curve as an example obtained for *n*_sh_ = 1.44, *n*_in_ = 1.363 and *t*_sh_ = 5 nm. The curve reproduces the data trend quite well. The larger spread in *C* as compared to the data for pure liposomes (see Fig. 4a) reveals detectable variations of *t*_sh_, *n*_sh_ or *n*_in_, which provide a gateway to quantitative studies of parasite secretion activity. For instance, if we assume *n*_sh_ = 1.44 and *t*_sh_ = 5 nm, as well as an effective RI of 1.6 for protein matter,^45^ we estimate the protein content of the EV inner solution to be about 10% *±* 3% as delineated by the olive curves.

### Human urine EVs

Urine is known to contain EVs, and scientists believe that these hold a great promise to serve as disease markers.^47,48^ Urine has been analyzed by NTA,^34,49^ yielding a unimodal vesicle size distribution in the range of 50 to 300 nm. Figure 4e shows, however, that although the iNTA size histogram also presents a continuous broad distribution, its 2D scatter plot can clearly resolve various sub-populations. Indeed, the observed bifurcation of the current data cloud starting at about *d* = 100 nm sheds a new light on particles present in urine. In Fig. 4f, we display isosurfaces for seven exemplary points marked A-G in Fig. 4e. All together, the scatter plot suggests a more complex situation than a simple categorization into two groups of exosomes and microvesicles.^6^ More analysis, in particular electron microscopy and fluorescent labeling, is needed to identify the urine constituents as EVs with full certainty, however the characteristic iNTA plot in Fig. 4e provides a strong basis for quantifying them.

## Discussion and outlook

In this work, we have introduced iNTA as an all-optical method that pushes the limits of sensitivity, precision and resolution in determining the size and refractive index of nanoparticle mixtures. We have demonstrated the power this technique by detecting nanoparticles that are more weakly scattering than previously reported, gaining insight into the hydration layer of colloidal gold nanoparticles, and deciphering complex nanoparticle species in various polydispersions. Moreover, we have shown that the current performance of iNTA is able to shed new light on medical diagnostics. In addition to its promise for boosting the analysis of samples extracted from the body, we see a particularly exciting application of iNTA to be in *in situ* characterization of extracellular vesicles,^37^ biomolecular condensates,^50^ viruses or other cellular nano-entities.

The method can also be improved further by several means, e.g., the use of shorter laser wavelength and higher laser power to increase the signal-to-noise ratio as well as the employment of particle confinement strategies^51^ to extend the measurement time. It is also important to note that iNTA can be readily combined with sensitive fluorescence measurements to extract additional information about the particles under study. These measures will give access to the high-resolution analysis of nanoparticles in a fast, precise and non-invasive fashion for a wide range of applications.^1–4,37^

## Acknowledgements

The authors are grateful to Alexey Shkarin the video acquisition software, Alexandra Schambony and Andreas Giessl for TEM measurements, Eduard Butzen for SEM measurements, Andreas Eigen (Halik lab, FAU) for support with DLS measurements, Alexander Zika (Gröhn lab, FAU) for support with NTA measurements and Felix Reichel (Guck lab, MPL) for support with viscometer measurements. We also thank Franziska Gröhn, Kai-Uwe Eckardt, Philipp Enghard, Rainer Böckmann, Mischa Bonn, David Albrecht, Mahdi Mazaheri and Kiarash Kasaian for helpful discussions. We are grateful to Michelle Küppers, Alexey Shkarin and Jennifer Lühr for a careful reading of the manuscript and insightful comments. We thank the Max Planck Society, and Alexander von Humboldt Foundation (fellowship for A.K.) for financial support.

## Author contributions

A.K., M.B., A.G., and V.S. conceived the experiments, M.B. performed the experiments, A.K., M.B., and A.G. analyzed the data. D.S. contributed materials. All authors discussed the results and commented on the manuscript. A.K., M.B., A.G., and V.S. wrote the paper. V.S. supervised the project.

